# Solid-state nanopore detection of partially denatured dsDNA with single-strand binding protein: a preliminary analysis

**DOI:** 10.1101/2024.01.28.577651

**Authors:** Nathan Howald, Alexander R. Klotz

## Abstract

In this work we investigate the use of a nanopore sensor to detect single-strand binding protein (SSB) attached to AT-rich denaturation bubbles in genomic double-stranded (ds) DNA. DNA from the *λ* bacteriophage was heated in the presence of E. coli SSB at temperatures predicted to open denaturation bubbles near the center of the molecule. A solid state nanopore sensor measured the ionic current as the DNA-SSB solution flowed through the pore, detecting blockades due to the translocation of biomolecules. Large current spikes were observed in the translocating DNA molecules, consistent with SSB binding. However, spikes were largely localized at either end of the DNA molecule, rather than at the predicted sites. We discuss the physico-chemical effects behind this disagreement and prospects for the future use of this technique for genomic mapping.

## I. INTRODUCTION

Rapid genomic mapping has the potential to revolutionize point-of-care diagnostics [1]. Unlike genetic sequencing, which provides base-level resolution, genomic mapping measures large-scale features of the genome and can provide information in much less time than sequencing, and can identify events such as gene duplications that may be missed by traditional sequencing technology [2]. One genomic mapping technique, pioneered by BioNano Genomics, confines and stretches DNA molecules in nanofluidic channels and looks at the distribution of fluorescent tags along the genome [3]. An improvement on the information density of tag-based nanochannel mapping can be achieved through denaturation mapping, which exploits the fact that AT-rich regions of the genome denature at lower temperatures than CG-rich regions (due to the fewer hydrogen bonds between AT), and that intercalating fluorescent dyes leave the melted sections of the molecule, allowing an optical barcode of the AT-richness along a molecule [4]. Denaturation barcoding is not a commercial technology, but it has been deployed in a hospital setting to identify antibiotic resistance in bacterial plasmids [5].

A nanopore sensor reads the ionic current as a fluid passes through a nanoscale pore in a membrane, and as biomolecules translocate through the pore they block the flow of ions, which is measurable as a drop in current. Nanopore sequencing, which measures the ionic current blocked by different bases along DNA molecules as they translocate through a nanoscale pore, has recently become a successful technology in the form of the Oxford MinIon [6]. The MinIon is the size of a USB key and uses a biological pore in a lipid membrane, builds a consensus sequence from several hundred parallel pores, and can sequence up to 420 bases per second. Solid state nanopores, in contrast, have not achieved base-level sensitivity but offer much faster translocation rates and the capability of processing more complex biomolecules, such as higher-order protein and RNA structures and DNA-protein complexes [7]. There has been recent success in developing genomic mapping schemes using solid state nanopores, including through the use of CRISPR [8] and nick-bound oligonucleotide tags [9].

In this work, we aim to combine the speed and large-scale information density of denaturation mapping with the low cost and miniaturization of nanopores. To do this, we will stabilize partially-denatured DNA using single-strand binding proteins (SSB) (Fig. 1a). SSB is used for regulating genome replication and forms complexes with 35 or 65 bases of single stranded DNA that wraps around the 4×4×7 nm protein [10]. Because the bound proteins have a larger cross section than bare DNA, they produce a strong signal in the current as they translocate, and are likely to be bound to regions rich in AT (Fig. 1bc). By tuning the parameters of the denaturation and binding protocols, we hope to develop an assay that can rapidly map the AT-rich regions of any DNA molecule without the need for sequence-specificity in the tags.

**FIG. 1:**
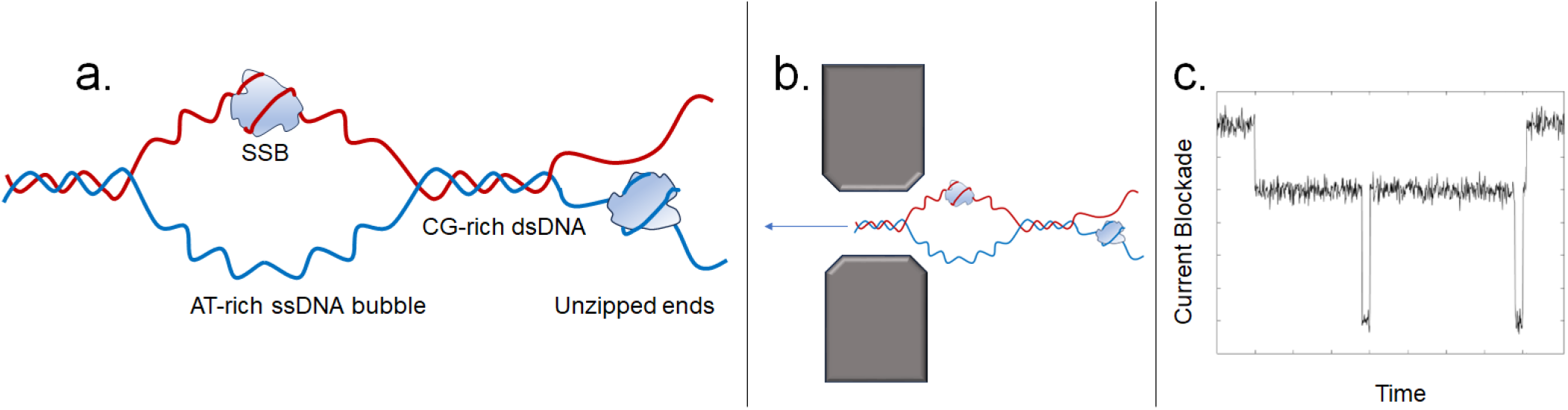
a. Schematic of a partially denatured DNA molecule with single strand binding proteins bound to an AT-rich denaturation bubble at the center, as well as to an unzipped end. b. Schematic of this DNA molecule translocating through a nanopore. c. Hypothetical current trace of such a molecule, in which the AT-rich regions and ends can be rapidly identified. The molecule pictured in this figure does not necessarily correspond to those used in this study, and the data in the plot is hypothetical.

Previously, it has been demonstrated that multiple SSB molecules will bind to a denaturation bubble formed by supercoiling. Complexes of SSB bound to single-stranded DNA have been detected using solid state nanopores, which have been used to explore binding kinetics [11] and show that the complex preferentially translocate protein-first [12]. To our knowledge, SSB has not previously been used for genomic mapping, nor has its binding to denaturation bubbles in dsDNA been investigated on a single-molecule level. In this work, we investigate genomic-length SSB-DNA complex as they translocate through a solid state nanopore as a preliminary investigation into the utility of this assay as a genomics technology.

## II. METHODS

We hybridized dsDNA from the *λ* bacteriophage (48,502 bp) with “extreme thermostable” E. coli SSB (both from New England Biolabs). In addition to being a readily available genome, *λ* has an AT-rich region at its center that would be easily identifiable in denaturation-based genomes, as has been seen previously [4]. One half of the molecule is significantly richer in AT-than the other, allowing directionality to be determined from SSB binding sites. Theoretical melting barcodes can be computed from the *λ* genetic sequence using the Bubbly Helix algorithm [13], which incorporates a thermodynamic model to determine whether a given part of the sequence will be open or closed. The location and size of the denaturation bubbles depends on the underlying sequence as well as the temperature and ionic strength. The predicted melting probability along *λ* DNA is shown in Fig. 2. The central denaturation bubble begins opening at around 77° C and expands with temperature. Other denaturation bubbles on the 3’ half of the molecule open at higher temperatures, and the ends are predicted to unzip significantly beginning at about 82° C. We chose 80° C and 2X TBE (63 mM at pH 8) to open a cluster of denaturation bubbles at the center of each dsDNA molecule, and a smaller one near one end of the chain. This is higher ionic strength than that used by Adamcik et al. to examine binding of SSB to twist-induced denaturation bubbles in dsDNA, but lower than that used by Japrung et al. [11] to bind SSB to natively single-stranded DNA. We note that under these conditions, unzipping at either end of the molecule is predicted to be unlikely, and any unzipped regions are likely smaller than denaturation bubbles in the interior. We also note that although SSB binds to 35 or 65 bases of ssDNA, the largest predicted denaturation bubble spans approximately 1500 bases.

**FIG. 2:**
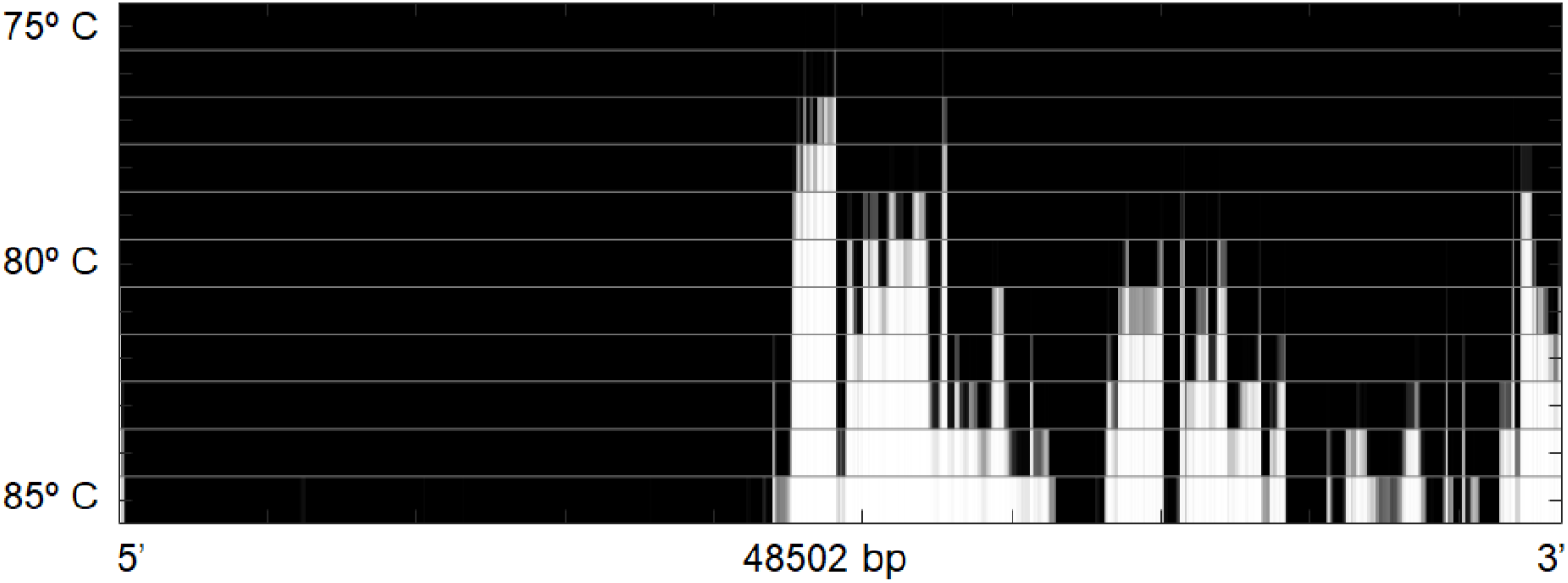
Theoretical melting probability along a *λ* DNA molecule from 75 to 85 degrees Celcius at 63 mM ionic strength. Brighter regions are more likely to be melted, and are the expected binding sites for SSB. As temperature increases, more binding sites become available. Most experiments in this study were at 80° C.

We followed similar procedures as those laid out by Japrung et al. using concentrations of DNA and SSB such that there are on average 4 SSBs per DNA. To do so we mixed 0.75 *µ*g of *λ* DNA and 6.5 ng of SSB (both from NEB) in 2X TBE and immersed it in a water bath at 80deg C for 15 minutes. After being removed from the heat and cooling to room temperature, the SSB-DNA solution was mixed with 4 M lithium chloride in a 1:3 ratio to yield a final salt concentration of slightly more than 3M. The complexation reaction took place at lower salt concentrations than the translocations to promote denaturation.

The nanopores were formed using controlled dielectric breakdown in the NanoCounter, a device from the now-defunct Ontera Inc. A microfluidic flow cell containing *cis* and *trans* channels separated by a silicon nitride membrane were loaded with 4M lithium chloride, and 10 volts were pulsed across the membrane until a pore across it. The pores typically reached a size of 30 nanometers, as determined by their conductivity. After fabrication, the SSB-DNA buffer was loaded into the *cis* channel. A bias of typically 100 mV was applied across the pore, and the ionic current through the pore was measured at a sampling rate of 125 kHz. Biomolecule translocations were identified by transient drops in the current.

## III. ANALYSIS AND RESULTS

Translocations can be characterized by their translocation time and mean and maximum current blockade. An example of an experimental run is visualized in scatter plots in Fig. 3. In both plots, two clusters are evident: one at times below half a millisecond with a broad range of currents, and one at longer times between 2 and 10 milliseconds, with the mean current having the characteristic inverse-time dependence of a constant electric-charge-deficit (ecd) hyperbola. The short-time cluster corresponds to unbound SSB molecules and clusters theoreof. E. coli SSB has an isoelectric point of 6.0 [14], making it translocate in the same direction as DNA in our experimental conditions. The longer-time cluster corresponds to DNA molecules, both with and without bound proteins. The maximum-current plot shows a clustering at around 100 and 200 pA, corresponding to the blockages caused by unfolded and folded unbound dsDNA. We applied a cutoff of 300 pA to separate unbound DNA from SSB-DNA complexes. This cutoff may lead to bound molecules categorized as unbound, but these false negatives are not major component of the data. The unbound molecules lie at the long-time-small-current end of the constant-ecd curve. There is a secondary long-time cluster with approximately twice the ecd, corresponding to simultaneous translocations. It is expected that bound SSB-DNA molecules have a greater average current blockade due to proteins blocking the pore; the shorter translocation compared to unbound DNA times may be interpreted as due to an effective shortening of the molecule as 65 bases of ssDNA span about 40 nm, which become wrapped around a protein 8 nm in length. The scatter plot of current maximum shows that the typical current magnitude of the bound spikes has a comparable distribution to unbound SSB.

**FIG. 3:**
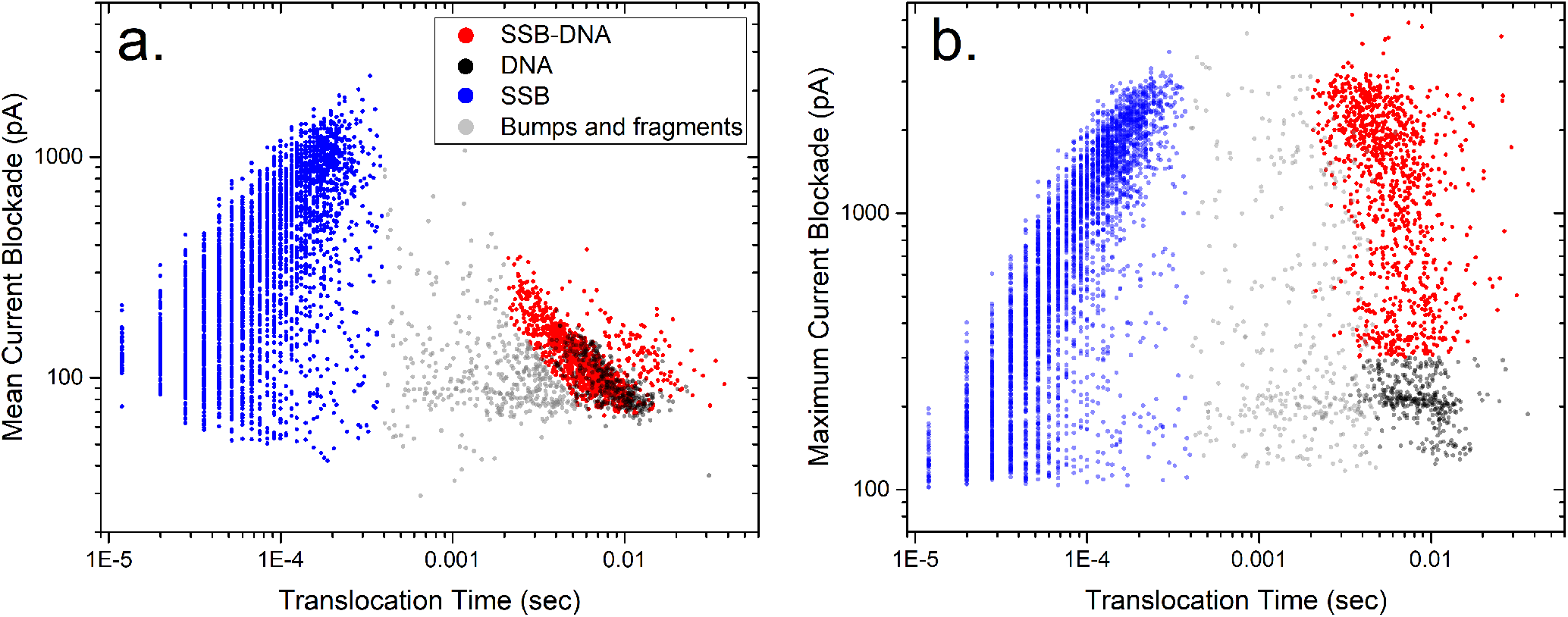
a. Scatter plot of the mean current blockade and duration of translocations from an experimental run through a 31 nm pore under a 100 mV bias. The cluster to the left is isolated SSB, the cluster to the right is DNA with and without bound proteins. In between lies bumps and fragmentary translocations. b. Scatter plot of the maximum current blockade and duration of translocations from the same experimental run.

The translocations with greater current deficits typically have a deep spike in their signal which we associate with bound SSB-DNA complexes. Example translocations can be seen in Fig. 4. A typical current depth for the SSB spike is on the order of 1000-2000 pA, which is comparable to the values seen for isolated SSB in Fig. 3b. Figure 5 shows histograms of the peak current amplitude for translocations from a DNA-only experiment, a DNA-SSB experiment, and the unbound SSB spikes. The peak current distribution for SSB-DNA has two peaks, a narrow one at low currents which matches well with the DNA-only experiment, and a broad one at high currents which matches the distribution of SSB.

**FIG. 4:**
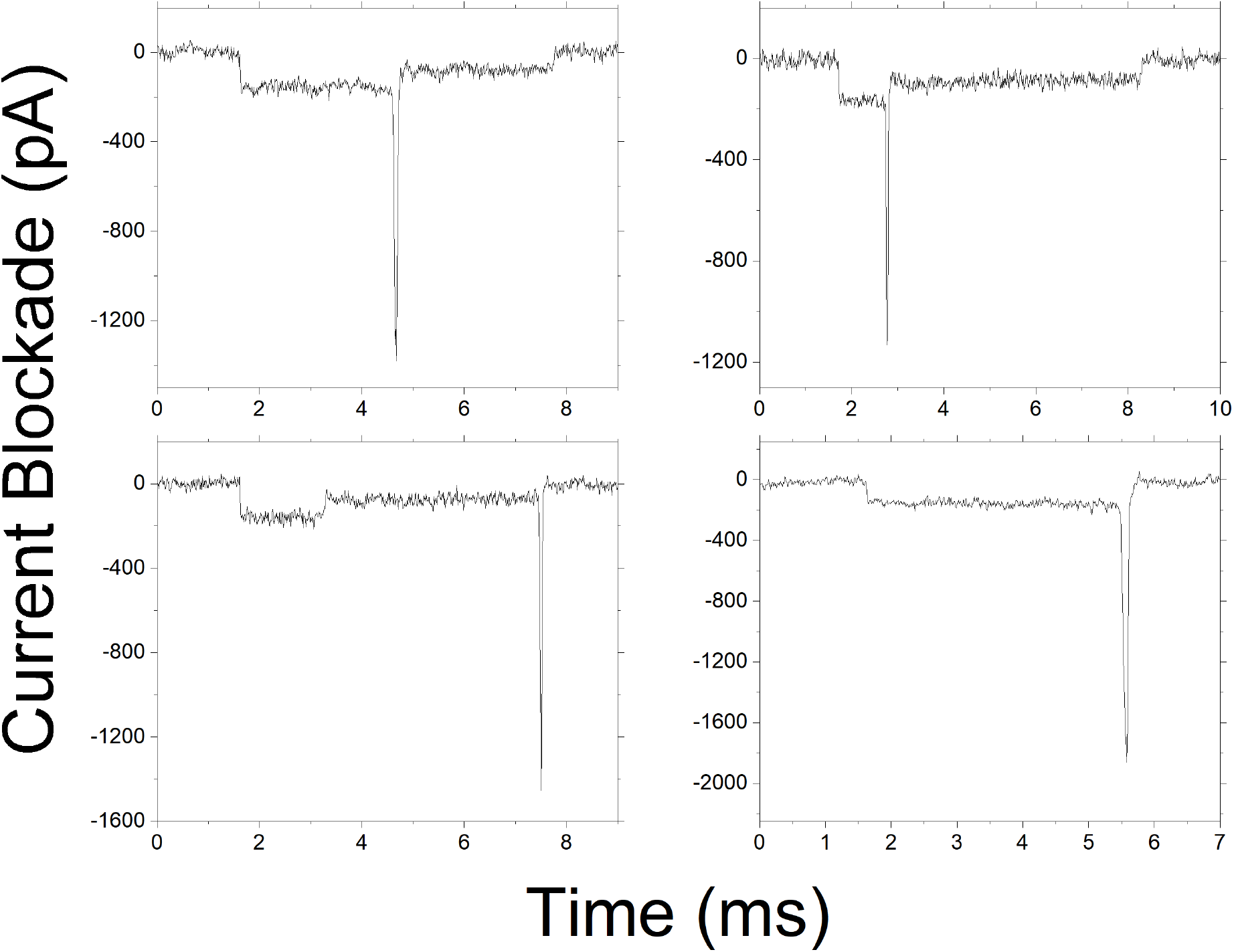
Four examples of the current blockades during translocations of a *λ* DNA molecule with SSB bound to the end. The top two traces have a fold at their leading end and have the SSB spike localized at the end of the fold. The bottom two have the SSB spike at their trailing end.

**FIG. 5:**
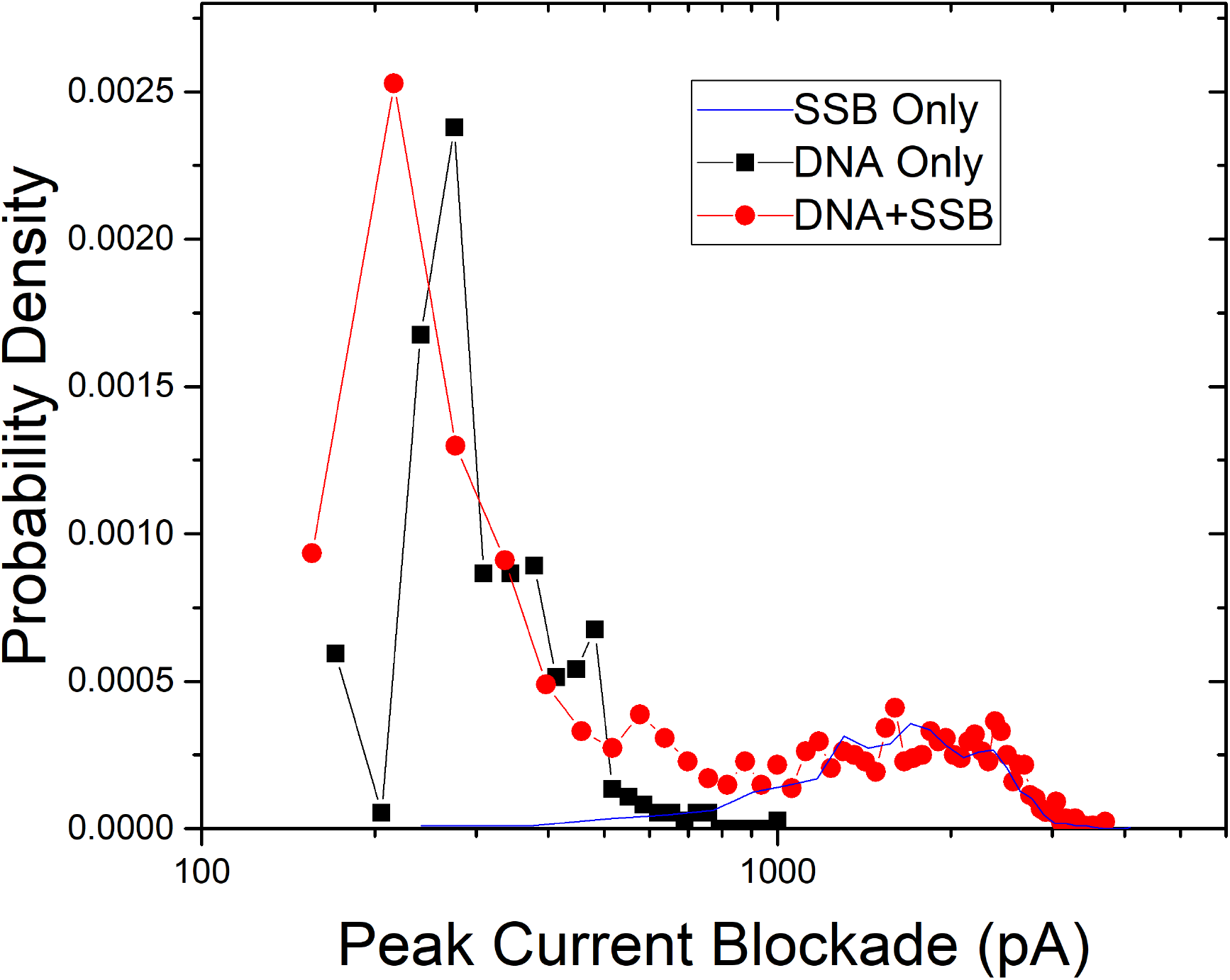
Histograms of peak current depth for SSB, DNA, and bound SSB-DNA complexes. The SSB+DNA histogram is bimodal, with one peak lying between the single- and double-fold translocation currents for unbound DNA, and the other tracing the distribution of unbound SSB molecules. Note that the SSB+DNA histogram is normalized, while the other two histograms are scaled to match the amplitude of the relevant part of the SSB+DNA set.

Examining individual translocations such as those in Fig. 4, the current spikes were overwhelmingly located near the ends of molecules. When the current minimum lay near the center of the molecule, it was most likely on a molecule with a fold, and was found at the edge of the fold. Although there may be two locations consistent with that spike, we interpret them as being located at molecule end. It is possible that the SSB is coincidentally localizing at a point on the molecule’s interior that happens to be where the fold ends. We consider this unlikely given how often we observe spikes near the straight ends of molecules. It is also possible that the SSB is binding an interior denaturation bubble to the molecule end into a lasso-like structure, but we do not believe that is the case. The agreement between the peak current distribution of SSB and SSB-DNA, as well as the non-uniform distribution of current spike locations, indicates that we are observing translocations of bound SSB-DNA complexes, and not simultaneous translocations of unbound SSB and DNA.

To categorize the location of SSB along the DNA, we found the time of each current minimum and divided it by the total translocation of the molecule to get an index between 0 and 1, a 0 corresponding to molecules which enter the pore SSB-first, and 1 corresponding to SSB-last. To handle folded molecules, we took molecules with indices between 0.1 and 0.9 and measured the mean current over 50 time points directly before and after the spike. If the mean currents before and after are close to a 2:1 ratio, it is interpreted as a fold (we used a threshold ratio of 1.5). If the fold entered the pore before the protein, we assign this an index of 0, and an index of 1 if the protein preceded the fold.

Figure 6 shows the SSB location index for 2563 lambda DNA translocations. The molecules are overwhelmingly located near either end of the molecule. Of the moelcules in the interior, there is a weak peak near the center, but this is not statistically significant. Although it appears that the SSB is equally likely to be found on either end, there is a bias that depends on the folding of the molecule. Straight molecules have a 2:1 bias towards translocating the pore with the SSB at the trailing end (an index close to 1), whereas folded molecules are overwhelming likely to translocate with the SSB-terminated fold entering first, as in the two current profiles in Fig 4. Molecules with the fold and SSB at the trailing end are essentially never observed. It is likely from this that the tagged ends are preventing from entering until the entire fold has translocated, which likely also explains the bias towards trailing-end proteins in straight molecules.

**FIG. 6:**
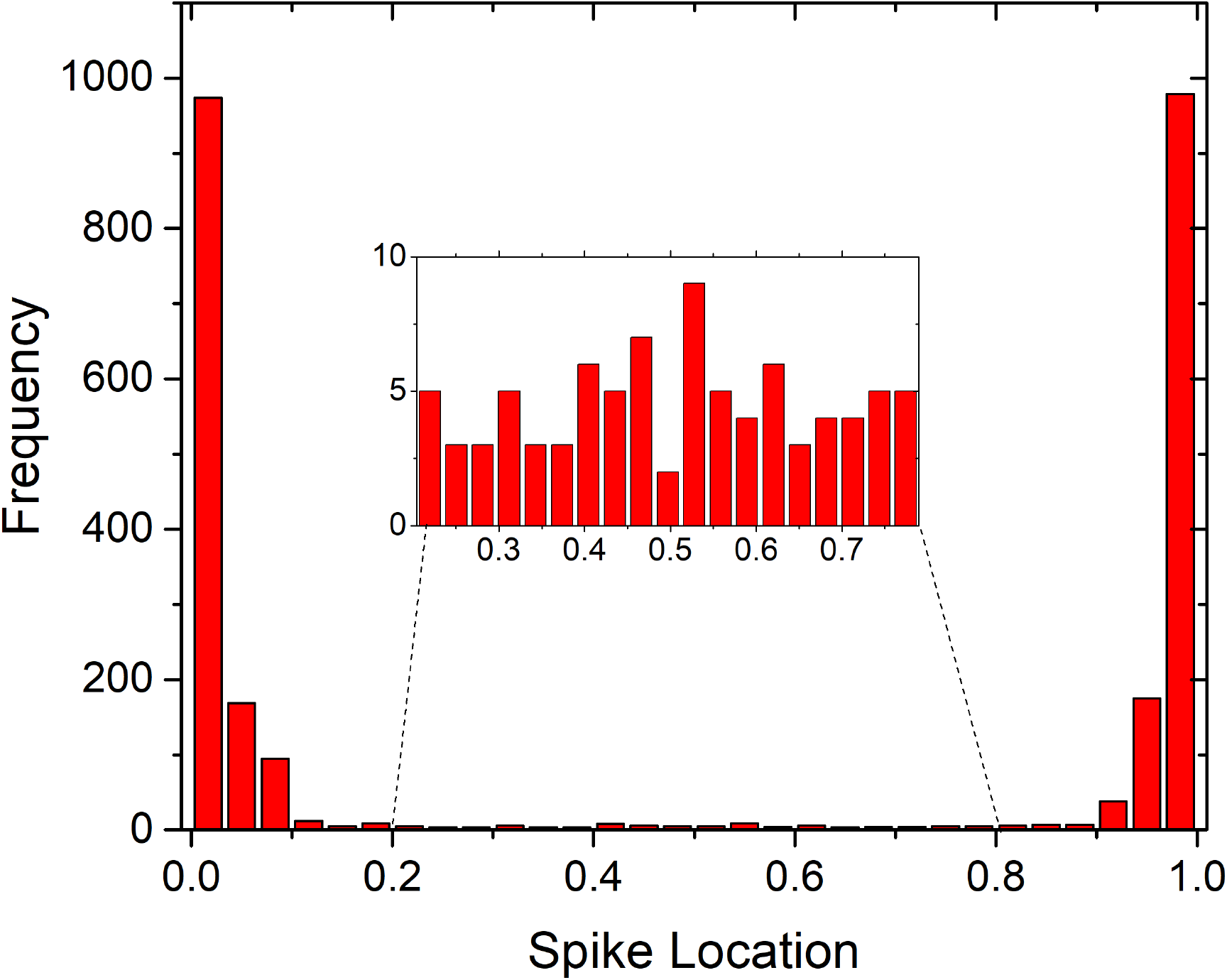
Histogram of measured binding locations of SSB along 2563 lambda DNA translocations, according to the categorization procedure defined in the text. They are clustered at the ends of the molecule. The inset shows the distribution in the interior of the molecule.

## IV. DISCUSSION

We anticipated that SSB would bind to denaturation bubbles at the center of the *λ* genome. This was not observed, instead SSB was almost universally bound to the chain ends. This is unexpected from the thermodynamic models used to predict the melting curves. There are several possible reasons for this discrepancy, relating to the validity of the melting models, differences in SSB binding kinetics to unzipped ends and internal bubbles, the stability of the SSB-DNA complexes within bubbles, and the particular features of *λ* DNA.

*λ* DNA has complementary single-stranded 12-base overhangs at each end of the genome. Although the smaller SSB-DNA complex forms with 35 base pairs, it is conceivable that our observations are consistent with the SSB binding to the overhangs. However, we have also performed experiments with T4 DNA (see Appendix), a longer viral genome without overhangs. While we do not have the same statistics as with our *λ* experiments, we also observe current spikes at the ends of the molecule (including at folds), suggesting the end-binding is not unique to *λ* DNA or its overhangs.

The DNA melting models are validated by calorimetric melting curves [15], and more recently through optical denaturation mapping in nanochannels [4]. The nanochannel experiments typically lower the melting temperatures significantly using formamide, which may change the thermodynamics. They also cannot directly observe unzipping at the ends, as it would simply reduce the apparent length of the molecule, which is already shrinking due to internal denaturation. The unzipping of ends in nanochannel-confined DNA can in the future be explored by combining YOYO-1 de-intercalation with sequence specific tags, for example those that bind to the overhangs of *λ* DNA.

If SSB binds to the ends at a much greater rate than bubbles, it may explain why bubble binding was not observed. Our binding assay typically took place over 15 minutes, longer than the successful binding reported by Unceuliac and Shuman after heating DNA for 5 minutes, rapidly cooling it on ice, and then mixing it with SSB [16]. Japrung et al. use a kinetics model to validate their observations of SSB binding to natively single-stranded DNA [11], but not to bubbles or unzips. The binding kinetics of SSB to denaturation bubbles have been investigated theoretically [17], although that work used clamped chain ends as boundary conditions and could not investigated the relative binding of unzips and bubbles. There is room for more investigation in this area, both calorimetric and computational.

There are competing factors in the desired salt concentrations for the binding assay, as the denaturation reaction and SSB binding have different salt dependences. At low salt concentrations, the 35-base binding mechanism is preferred, while at high salt concentrations, the 65-base mechanism is preferred. Our experiments took place in an intermediate regime. Experiments from Hatch et al. [18] use force-induced unzipping to examine the stability of DNA-SSB complexes and find that at higher salt concentrations, the complexes are not stable and the DNA re-zips when the force is removed, whereas at lower salt concentrations, the SSB can prevent re-zipping when the force is removed. The proposed mechanism is that the 35-base structure is cooperative and essentially forms filaments along the ssDNA (as seen by Marshall et al [12]) whereas at higher salts the 65-base structure is not cooperative, and helices may reform on either side of the SSB complexes, possibly forcing them off the molecule.

The binding failure of SSB in the weak force regime may explain why do not observe it binding to the molecule’s interior. It does not, however, explain why we observe them binding to the unzipped ends. This highlights a lack in our understanding of the difference between bubbles and unzips, and the binding mechanisms of proteins to them. It also does encourage the development of this assay as a genomic mapping technology. However, future experiments exploring a broader region of the parameter space (particularly with lower ionic strengths where the 35-base structure dominates) as well as using the T4 Gene 32 protein instead of E. Coli SSB (which is known to be stabler at low forces [19], may prove a fruitful avenue of investigation.

## V. ACKNOWLEDGEMENTS

We are grateful to Kevin Freedman, Wayne Yang, and Tobias Ambjörnsson for helpful discussions. This work was supported by the National Science Foundation (Grant number 2105113) and a CSUPERB New Investigator Grant.

## VI. AUTHOR DECLARATION

The authors declare no financial conflicts of interest but are required to publish papers to advance their careers.

## VII. APPENDIX

Figure 7 shows four translocations of T4 DNA with SSB bound at or near the end of the molecule. These translo- cations were measured at a 200 mV bias voltage, compared to the 100 mV *λ* DNA experiments. We did not observe a sufficient number of T4 translocations to discuss statistics, but we do observe frequent localization of the spikes at the ends of the molecules.

**FIG. 7:**
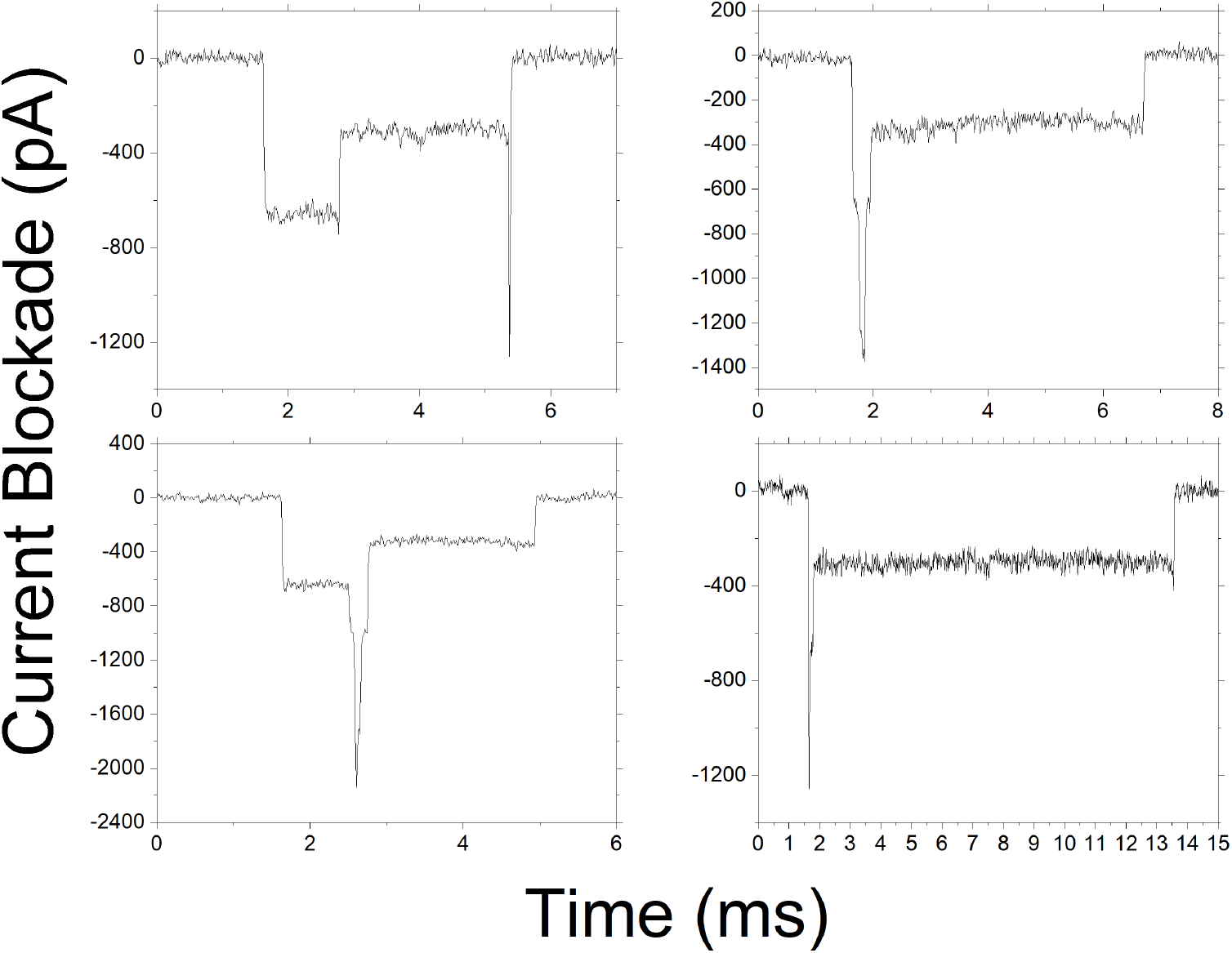
Four examples of the current blockades during translocations of a *T* 4 DNA molecule with SSB bound to the end.

## Notes

### Competing Interest Statement

The authors have declared no competing interest.

## References

[1] Kewal K Jain and Kewal K Jain. Molecular diagnostics in personalized medicine. Textbook of Personalized Medicine, pages 39–101, 2021.

[2] Paul Dremsek, Thomas Schwarz, Beatrix Weil, Alina Malashka, Franco Laccone, and Jürgen Neesen. Optical genome mapping in routine human genetic diagnostics—its advantages and limitations. Genes, 12(12):1958, 2021.

[3] Sven Bocklandt, Alex Hastie, and Han Cao. Bionano genome mapping: high-throughput, ultra-long molecule genome analysis system for precision genome assembly and haploid-resolved structural variation discovery. Single Molecule and Single Cell Sequencing, pages 97–118, 2019.

[4] Walter Reisner, Niels B Larsen, Asli Silahtaroglu, Anders Kristensen, Niels Tommerup, Jonas O Tegenfeldt, and Henrik Flyvbjerg. Single-molecule denaturation mapping of dna in nanofluidic channels. Proceedings of the National Academy of Sciences, 107(30):13294–13299, 2010.

[5] Vilhelm Muller, Nahid Karami, Lena K Nyberg, Christoffer Pichler, Paola C Torche Pedreschi, Saair Quaderi, Joachim Fritzsche, Tobias Ambjornsson, Christina Åhrén, and Fredrik Westerlund. Rapid tracing of resistance plasmids in a nosocomial outbreak using optical dna mapping. ACS infectious diseases, 2(5):322–328, 2016.

[6] Miten Jain, Hugh E Olsen, Benedict Paten, and Mark Akeson. The oxford nanopore minion: delivery of nanopore sequencing to the genomics community. Genome biology, 17:1–11, 2016.

[7] Chalmers Chau, Fabio Marcuccio, Dimitrios Soulias, Martin Andrew Edwards, Andrew Tuplin, Sheena E Radford, Eric Hewitt, and Paolo Actis. Probing rna conformations using a polymer–electrolyte solid-state nanopore. ACS nano, 16(12):20075–20085, 2022.

[8] Wayne Yang, Laura Restrepo-Pérez, Michel Bengtson, Stephanie J Heerema, Anthony Birnie, Jaco Van Der Torre, and Cees Dekker. Detection of crispr-dcas9 on dna with solid-state nanopores. Nano letters, 18(10):6469–6474, 2018.

[9] Arthur Rand, Philip Zimny, Roland Nagel, Chaitra Telang, Justin Mollison, Aaron Bruns, Emily Leff, Walter W Reisner, and William B Dunbar. Electronic mapping of a bacterial genome with dual solid-state nanopores and active single-molecule control. ACS nano, 16(4):5258–5273, 2022.

[10] Wlodzimierz Bujalowski and Timothy M Lohman. Escherichia coli single-strand binding protein forms multiple, distinct complexes with single-stranded dna. Biochemistry, 25(24):7799–7802, 1986.

[11] Deanpen Japrung, Azadeh Bahrami, Achim Nadzeyka, Lloyd Peto, Sven Bauerdick, Joshua B Edel, and Tim Albrecht. Ssb binding to single-stranded dna probed using solid-state nanopore sensors. The journal of physical chemistry B, 118(40):11605–11612, 2014.

[12] Michael M Marshall, Jan Ruzicka, Osama K Zahid, Vincent C Henrich, Ethan W Taylor, and Adam R Hall. Nanopore analysis of single-stranded binding protein interactions with dna. Langmuir, 31(15):4582–4588, 2015.

[13] Eivind Tøstesen, Geir Ivar Jerstad, and Eivind Hovig. Stitchprofiles. uio. no: analysis of partly melted dna conformations using stitch profiles. Nucleic acids research, 33(suppl 2):W573–W576, 2005.

[14] Timothy M Lohman and Marilyn E Ferrari. Escherichia coli single-stranded dna-binding protein: multiple dna-binding modes and cooperativities. Annual review of biochemistry, 63(1):527–570, 1994.

[15] Ralf Blossey and Enrico Carlon. Reparametrizing the loop entropy weights: effect on dna melting curves. Physical Review E, 68(6):061911, 2003.

[16] Mihaela-Carmen Unciuleac and Stewart Shuman. Double strand break unwinding and resection by the mycobacterial helicase-nuclease adnab in the presence of single strand dna-binding protein (ssb). Journal of Biological Chemistry, 285(45):34319–34329, 2010.

[17] Tobias Ambjornsson and Ralf Metzler. Blinking statistics of a molecular beacon triggered by end-denaturation of dna. Journal of Physics: Condensed Matter, 17(49):S4305, 2005.

[18] K Hatch, Claudia Danilowicz, V Coljee, and Mara Prentiss. Measurement of the salt-dependent stabilization of partially open dna by escherichia coli ssb protein. Nucleic acids research, 36(1):294–299, 2008.

[19] Kiran Pant, Richard L Karpel, Ioulia Rouzina, and Mark C Williams. Salt dependent binding of t4 gene 32 protein to single and double-stranded dna: single molecule force spectroscopy measurements. Journal of molecular biology, 349(2):317–330, 2005.

